# Tunneling CARs: Increasing CAR T tumor infiltration through the overexpression of *MMP7* and *SPP1*

**DOI:** 10.1101/2025.01.24.634731

**Authors:** Stacey N. Van Pelt, Mark White, Candise Tat, Devyn Hooper, Lindsay Talbot, Rohan Fernandes, Cliona Rooney, Bilal Omer

**Affiliations:** Graduate Program in Immunology and Microbiology, Baylor College of Medicine, Houston, TX 77030; Center for Cell and Gene Therapy, Baylor College of Medicine, Texas Children’s Hospital, and Houston Methodist Hospital, Houston, TX 77030; Department of Bone Marrow Transplantation and Cellular Therapy, St. Jude Children’s Research Hospital, Memphis, TN 38105; Department of Surgery, St. Jude Children’s Research Hospital, Memphis, TN 38105; The Institute for Biomedical Sciences, School of Medicine and Health Sciences, George Washington University, Washington, DC 20052; The George Washington Cancer Center, The George Washington University, Washington, DC 20052; Department of Medicine, School of Medicine and Health Sciences, George Washington University, Washington, DC 20052; Graduate Program in Translational Biology and Molecular Medicine, Baylor College of Medicine, Houston, TX; Dan L Duncan Comprehensive Cancer Center, Baylor College of Medicine, Houston, TX 77030; Department of Pediatrics, Section of Hematology-Oncology, Baylor College of Medicine, Houston, TX 77030; Department of Pathology-Immunology, Baylor College of Medicine, Houston, TX 77030

## Abstract

Chimeric antigen receptor T cell (CART) therapy has demonstrated remarkable efficacy in hematologic malignancies but has struggled to achieve comparable success in solid tumors. A key obstacle is the extracellular matrix (ECM) in solid tumors, which significantly impedes CART cell infiltration. In clinical trials, neuroblastoma (NB) has shown responsiveness to GD2-directed CART therapy, however, the failure of GD2.CARTs to effectively clear bulky disease - characterized by dense ECM - highlights the critical challenge of infiltration.

In this study, we demonstrate that GD2.CARTs exhibit a unique infiltration-restriction compared to other CARTs and endogenous T cells. A separate analysis of clinical datasets identified *MMP7* and *SPP1* (OPN) as candidate genes to improve the infiltration of GD2.CARTs as these were upregulated in tumor-infiltrating leukocytes.

MMP-7 and OPN overexpression enhanced CART extravasation (*p < .001*) and interstitial movement (*p < .05*) in ECM-dense environments *in vitro*. Overexpression of either OPN (*p < .0001*) or MMP-7 (*p < .001*) improved tumor infiltration in a xenograft model of NB. This resulted in improved tumor control (94% reduction in tumor burden, *p < .05*) and a survival extension in OPN-GD2.CART treated mice compared to unmodified GD2.CARTs (median of 148 days, *p < .05*). OPN overexpression did not increase off-target infiltration into healthy tissues or promote tumor metastasis, highlighting its potential for safe therapeutic application. Our study provides a framework for further exploration of gene modifications to improve CART infiltration and efficacy in solid tumors and identifies *SPP1* as a candidate gene to improve GD2.CART treatment of bulky tumors.

## INTRODUCTION

Chimeric antigen receptor T cell (CAR T) therapy has produced remarkable response rates in hematologic malignancies but complete responses in the solid tumor setting remain elusive^1–11^. A common observation from solid tumor trials is that responses correlate with tumor burden, where patients with higher tumor burden or bulky disease had poorer outcomes than those with smaller, more diffuse tumors^12–14^. One reason for this is the low infiltration of CAR T cells into tumors due to the exclusionary nature of the tumor-specific extracellular matrix (ECM), and basement membrane (BM) which encapsulate the tumor^15–17^.

Tumor encapsulation occurs when cancer associated fibroblasts (CAFs) recruited to the tumor secrete large volumes of ECM fibers, while simultaneously secreting enzymes that crosslink these fibers into a dense, rigid structure^18,19^. Unlike the ECM and BM of healthy tissues, which have a loose basket-weave pattern that T cells easily traverse, tumor-specific matrices are stiff and characterized by small pores, leading to T cell exclusion^19–21^.

T cells traveling to tumor sites through the circulation use rolling and firm adhesion to bind to the endothelium of the vasculature, then extravasate through the BM into the interstitial space and use adhesion molecule-mediated cell movement and cytoskeletal rearrangements to move through the tumor-specific ECM and into the tumor core^22–24^. Both barriers encountered by T cells in this process, the BM, composed mainly of collagen type IV, and the encapsulating tumor-specific ECM, composed of mainly collagen type I, must be enzymatically degraded by T cells via distinct matrix metalloproteinases (MMPs) to provide space for T cell movement^25–26^.

These barriers, while critical for regulating normal tissue integrity, pose significant challenges for immune cells targeting solid tumors. In the case of neuroblastoma (NB)—a cancer arising from neural crest-derived cells and the most common extracranial tumor in children—these challenges are further compounded by the tumor’s dense, rigid ECM. A defining characteristic of treatment-resistant, ultra-high-risk NB is a dense, rigid ECM^28^, that promotes tumor cell proliferation and inhibits the infiltration of T cells into tumor cores^29^.

Neuroblastoma highly expresses the tumor associated disialoganglioside GD2^27^. Unlike most other solid tumor-targeting CARs, complete responses have been achieved with GD2.CAR T cells, however, results have been modest in patients with high disease burden and larger tumors^12–14^.

There are few reports on the infiltration properties of CAR T cells or attempts to improve their infiltration potential^30,31^. Here, using publicly available datasets from patient samples derived from four different solid tumor types, we systematically identified key infiltration-associated genes, *MMP7, MMP14,* and *SPP1* (OPN), that were upregulated in tumor infiltrating leukocytes compared to those that were tumor-adjacent or circulating. We overexpressed these genes in GD2.CAR T cells and showed that *MMP7* and *SPP1* expression increased their infiltration in a preclinical model of NB, and that the overexpression of *SPP1* increased tumor clearance and survival.

## RESULTS

To visualize the distribution of endogenous T cells and ECM within solid tumors, we stained NB, breast cancer (BC) and pancreatic ductal adenocarcinoma (PDAC) samples for CD3 and collagen. We observed areas of bulk tumor tissue encapsulated by dense layers of collagen (tumor nests) within all three tumor types (**Fig 1A**). Rather than associating with bulk tumor tissue, endogenous T cells were typically localized within dense ECM surrounding these nests (**Fig 1A**). This distribution contrasted with T cell localization within healthy tonsil tissue, where T cells associated equally with dense ECM and bulk lymphoid tissue, suggesting tumor-specific ECM presents unique challenges for T cell movement.

**Figure 1.**
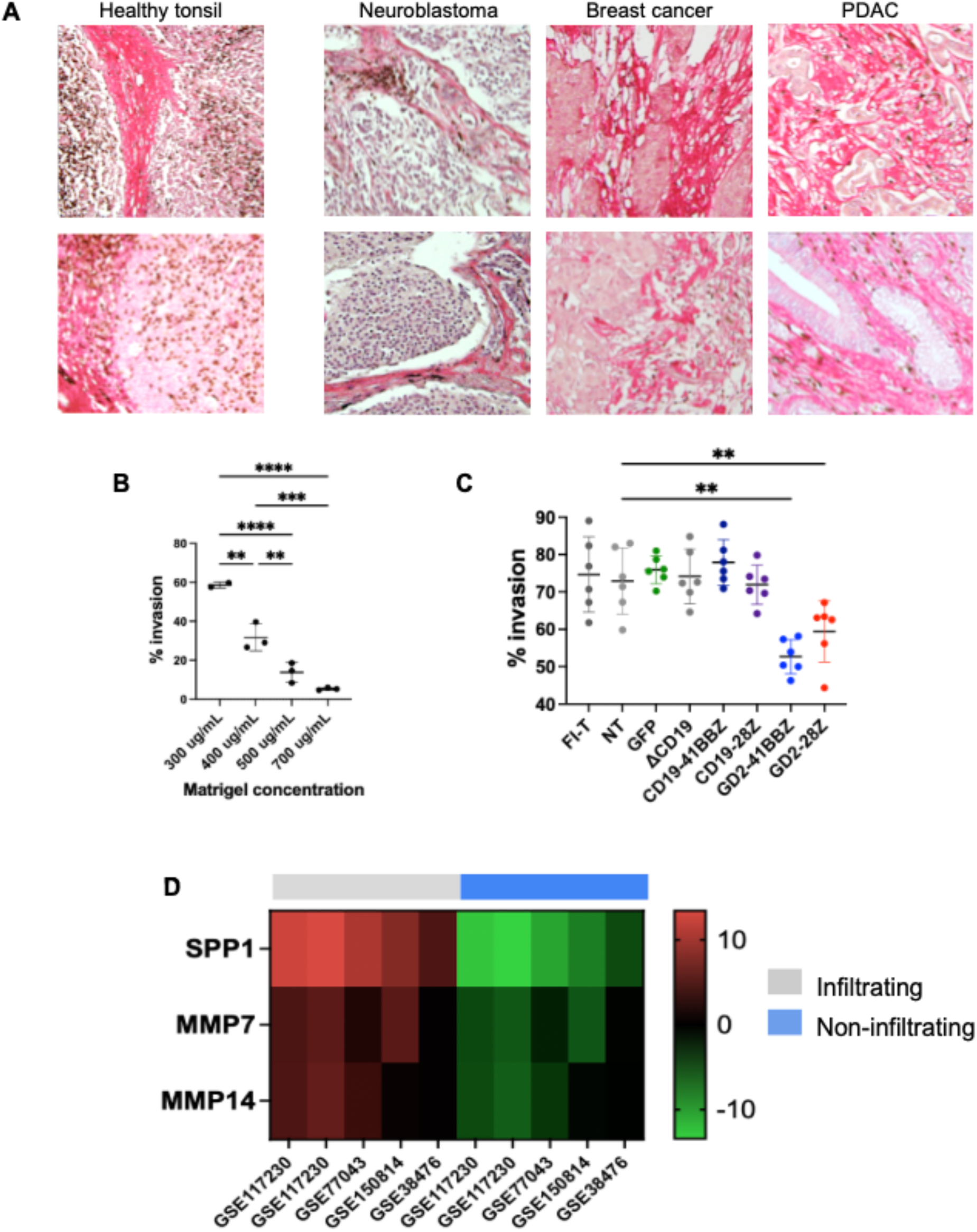
GD2.CAR T cells are infiltration-restricted through tumor-specific ECM. **A.** Patient biopsies of NB, BC and PDAC stained for collagen (red) and CD3 (brown) showing the aggregation of endogenous T cells in dense collagen surrounding tumor nests. **B.** Transwell invasion assay comparing infiltration of ATCs through increasing concentrations of Matrigel. T cell infiltration was highest (59% ± 1%) at the lowest concentration of Matrigel (300 mg/mL), and was most restricted at the highest ECM concentration of 700 mg/mL (5% ± 1%, *n = 2-3, p < .0001*). **C.** Transwell invasion assay comparing infiltration of freshly isolated T cells (FI-T), nontransduced T cells (NT), and T cells expressing CD19-41BBZ, CD19-28Z, GD2-41BBZ, or GD2-28Z CARs. T cells transduced with GFP or ΔCD19 were used as controls. Both GD2-41BBZ (53% ± 5%) and GD2-28Z (59% ± 8%) cells were infiltration-restricted compared to NT T cells (73% ± 9%, *n = 6, p < .01*). Data summaries are the mean ± the standard deviation, one-way ANOVA with Tukey’s correction (B) or Dunnett’s correction (C). **D.** Heat map of RNA expression in patient samples for candidate genes *MMP7, MMP14,* and *SPP1* in tumor-infiltrating and non-infiltrating leukocyte populations. Data was gathered from NCBI’s GEO database. *\*p < .05, **p < .01, ***p < .001, ****p < .0001*.

To understand how dense ECM affects the movement of *ex vivo* expanded T cells, we used transwell invasion assays to measure infiltration through increasing concentrations of Matrigel, which is composed mostly of collagen type IV, and models the basement membrane. Our results showed that at low concentrations of Matrigel, more than half of the cells were able to infiltrate (59% ± 2%, mean ± SD), but when concentrations were increased to levels closer to that of the tumor-specific ECM, infiltration sharply dropped to just 5% (± 1%, *p < .0001*) (**Fig 1B**) suggesting ECM density contributes to T cell aggregation, as expected.

Previous studies have shown that *ex vivo* expanded T cells poorly infiltrate through ECM compared to freshly isolated T cells (FI-T), but how infiltration is affected by CAR expression is currently unknown^1^. Further, unique characteristics of CARs, such as tonic signaling or choice of costimulatory domain may impact CAR T cell movement. To evaluate differences in infiltration between tonically-signaling (GD2.CAR) and non-tonically signaling (CD19.CAR) CAR T cells and the effects of CD28 and 4-1BB costimulatory domains, we compared the infiltration of FI-T, nontransduced activated T cells (NT), CD19-41BBZ, CD19-28Z, GD2-41BBZ, and GD2-28Z CAR T cells using a transwell invasion assay. T cells transduced with GFP or ΔCD19 were used as controls.

Although there were no differences in infiltration between CAR T cells expressing 4-1BB or CD28 costimulatory domains, or non-tonically signaling CARs (CD19-41BBZ, CD19-28Z) and FI-T, NT, GFP or ΔCD19 controls, we did observe a significant drop in the infiltration capacity of CAR T cells expressing the tonically-signaling GD2.CAR (GD2-41BBZ and GD2-28Z). These data indicate that endogenous T cells poorly infiltrate solid tumors, and instead aggregate in dense ECM surrounding tumor nests, and that GD2.CAR T cells are even more severely restricted. Thus, gene- modifications to enhance the interstitial movement of GD2.CAR T cells and their infiltration through ECM may improve their limited clinical efficacy against solid tumors.

### MMP7, MMP14, and SPP1 are upregulated in tumor infiltrating leukocytes

To identify infiltration-associated candidate genes, we analyzed four publicly available datasets (GSE117230, GSE77043, GSE183349, GSE38476) from the National Center for Biotechnology Information’s (NCBI) Gene Expression Omnibus (GEO) database comparing gene expression in infiltrating leukocytes with non-infiltrating or circulating leukocytes from patient samples in various tumor types. We first identified the top 50 differentially expressed genes in the infiltrating populations for each study, then cross-referenced those genes to identify candidates that were upregulated in more than one study. That analysis identified three candidate genes: *MMP7, MMP14*, and *SPP1* that were upregulated in infiltrating leukocytes (**Fig 1D**)*. MMP7* encodes a secreted protease that degrades collagen type IV, the main component of the BM. *MMP14* encodes a membrane bound protease that degrades collagen type I, the main component of the tumor-specific ECM, which encapsulates the tumor and surrounds tumor nests^41^. *SPP1* encodes osteopontin (OPN), a secreted protein that binds to ECM fibers, including collagens type I and type IV^33^, and acts as a ligand for integrins and adhesion molecules that mediate T cell extravasation and interstitial movement^34,35^. An intracellular isoform of OPN may also impact cell movement by anchoring the actin cytoskeleton to cytoplasmic domains of adhesion molecules that connect to the ECM^36^. Our analysis showed that *SPP1* was highly upregulated in infiltrating cells across all four studies, while *MMP7* and *MMP14* were each upregulated in two studies. Thus, we hypothesized that overexpressing these candidate genes in CAR T cells would increase their infiltration into solid tumors.

To screen our candidate genes, we transduced primary T cells with retroviral vectors (**Supplementary** Fig 1A) encoding *OPN* isoforms (*OPNa*, *OPNb*, *OPNc,* which are secreted isoforms, and the intracellular isoform *iOPN*), *MMP7* or *MMP14*, then used transwell invasion assays to measure how the expression of these genes affected infiltration *in vitro*. Results showed that the expression of *OPNb* (hereafter OPN) and *MMP7* increased T cell infiltration by a mean of 1.8- and 2.4-fold, respectively (**Supplementary** Fig 1C-D) over nontransduced (NT) T cells. In contrast, the expression of *MMP14* restricted infiltration and was therefore excluded from subsequent testing in GD2.CAR T cells (**Supplementary** Fig 1E).

### Tunneling GD2.CARs maintain key functions of GD2.CAR T cells

To generate “tunneling” GD2.CARs, we sequentially transduced activated T cells (ATCs) with the GD2-28Z CAR (GD2.CAR, **Fig 2A**) and then MMP-7 (MMP7-GD2.CAR) or OPN (OPN-GD2.CAR). NT T cells and single-transduced GD2.CAR T cells were used as controls. T cells were expanded and their transduction, phenotype, transgene expression and cytotoxicity were assessed. The transduction efficiency of the GD2.CAR alone was 88% ± 5% (mean ± SD), *n* = 4. Double- transduction efficiency, measured as the percentage of cells positive for both the GD2.CAR and the transgene marker, was 83**%** ± 6% for MMP7-GD2.CARs, and 71% ± 5% for OPN-GD2.CARs (**Fig 2B-C**). There were no significant differences in the expansion of GD2.CAR T cells, compared to MMP7-GD2.CAR or OPN-GD2.CAR T cells (**Fig 2E**).

**Figure 2.**
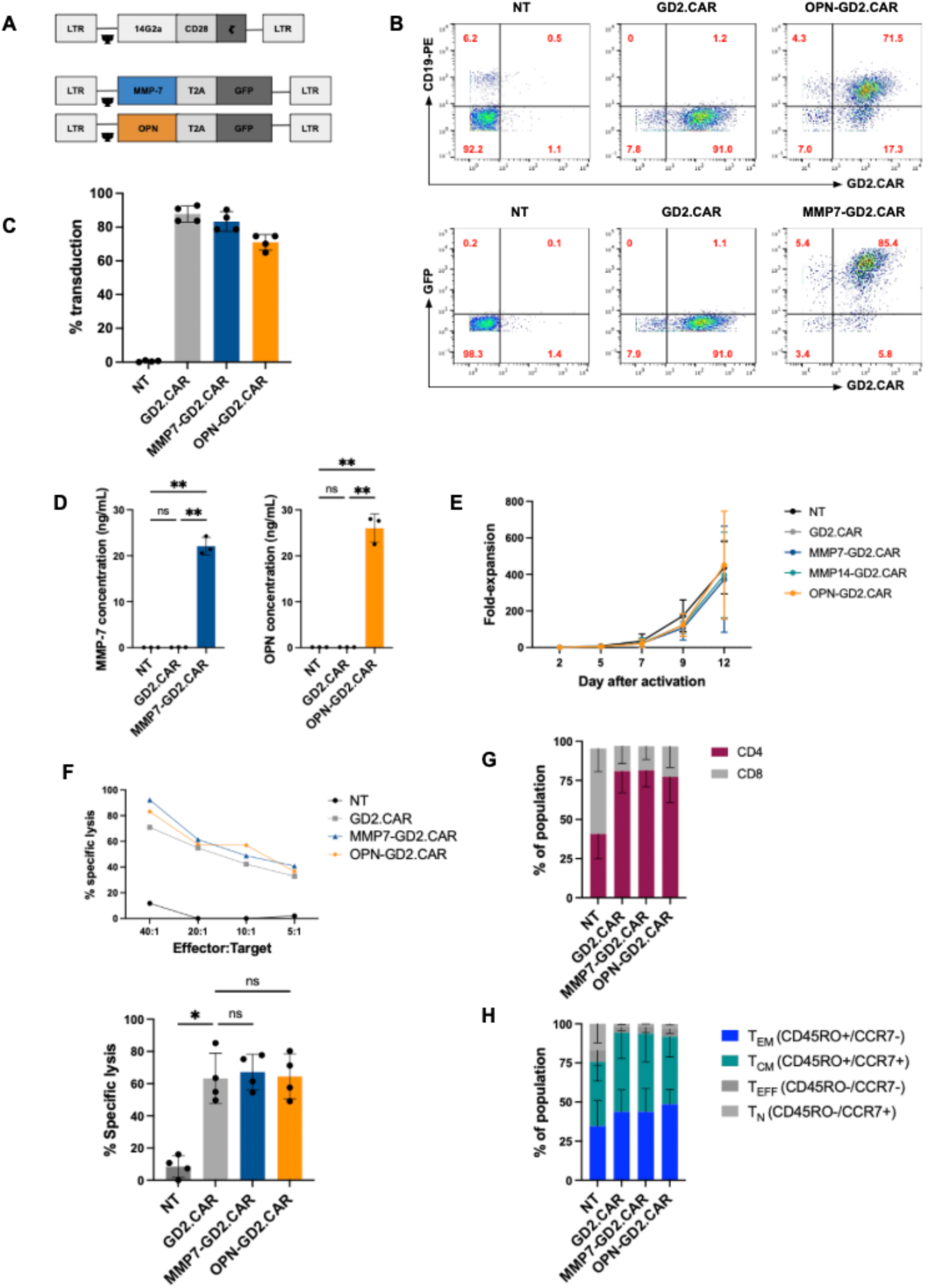
The expression of MMP-7 or OPN does not affect the expansion, phenotype or cytotoxicity of GD2.CAR T cells. **A.** Schematic of retroviral vectors of the GD2.CAR and candidate genes. **B.** Flow cytometry plots from a representative donor showing transduction efficiency of single-transduced GD2.CAR T cells, and double-transduced MMP7-GD2.CAR and OPN-GD2.CAR T cells. NT T cells were used as a control. **C.** Transduction efficiency of unmodified GD2.CAR T cells and modified GD2.CAR T cells from 4 donors. **D.** Secretion of MMP-7 (left) or OPN (right) by modified GD2.CAR T cells compared to unmodified GD2.CAR T cells measured by ELISA ten days post-activation (mean, SD, *n = 4*, one-way ANOVA with Tukey’s correction). **E.** Expansion of modified GD2.CAR T cells over 12 days after activation compared to unmodified GD2.CAR T cells (*n = 4*). **F.** Cytotoxicity of modified and unmodified GD2.CAR T cells at various effector:target ratios from a representative donor (top) and summary of data from all donors (bottom) at an effector:target ratio of 20:1 (mean, SD, *n = 4*, mixed effects analysis with Dunnett’s correction). **G.** Percentage of CD4 and CD8 T cells in the total population (mean, SD, *n = 4*). **H.** Phenotypic ratios of modified GD2.CAR T cells and unmodified GD2.CAR T cells measured eight days after activation (mean, SD, *n = 4*). *\* p < .05, **p < .01*.

To confirm the expression of these transgenes, we measured MMP-7 secretion by MMP7- GD2.CAR (22.1 ng/mL ± 1.9, *n = 3*), GD2.CAR (0.9 ng/mL) and NT T cells (0.1 ng/mL), and OPN secretion by OPN-GD2.CAR (26 ng/mL ± 3.1), GD2.CAR (0.1ng/mL), and NT T cells (0.1 ng/mL) by ELISA 8 days post-transduction (**Fig. 2D**). To ensure transgene expression did not disrupt the cytotoxic function of the GD2.CAR, we evaluated cytotoxicity using a short-term chromium release assay and found no significant differences in cytotoxicity between GD2.CAR (63% ± 7%, *n = 4*, effector to target ratio of 20:1), MMP7-GD2.CAR (67% ± 11%, *n = 4)* or OPN-GD2.CAR (64% ± 14%, *n = 4*) T cells (**Fig. 2F**). To determine the effects of MMP-7 or OPN expression on T cell phenotype we evaluated CD4:CD8 ratios, and effector:memory phenotypes, defined as effector memory (T_EM_; CD45RO+/CCR7-), central memory (T_CM_; CD45RO+/CCR7+), effector (T_EFF_; CD45RO-/CCR7-), and naïve (CD45RO-/CCR7+) T cells. We did not detect any significant differences between MMP7-GD2.CAR, OPN-GD2.CAR and GD2.CAR T cells in regards to their CD4:CD8 ratio (**Fig 2G**) or effector:memory subtypes (**Fig 2H**).

### Tunneling CARs exhibit superior infiltration in vitro

To evaluate how MMP-7 and OPN overexpression affected GD2.CAR T cell movement and invasion properties *in vitro*, we modeled the extravasation (defined here as movement through the BM) and infiltration of T cells using a Halo infiltration assay. GFP-transduced CAR T cells (green) were resuspended in high-concentration Matrigel (BM) and plated in a halo on the perimeter of a well, while tumor targets (red) were resuspended in high-concentration collagen type I and plated as a droplet in the center of the well (**Fig 3A**). T cell extravasation out of the BM, and infiltration into the tumor droplet, were then visualized over time (3-4 days) using live cell imaging. Extravasation was measured as the total T cell area per well at early time points of the assay (t = 6-12h), and infiltration was measured as the total area per well where T cells overlapped the tumor droplet. Our results showed that at early time points (12 hours) MMP7- GD2.CAR (1.25e7 um^2^ ± 9.18e5 um^2^, mean, SD, *n = 4, p < .001*), and OPN-GD2.CAR T cells (8.52e6 um^2^ ± 7.70e5 um^2^, *p < .001*) exhibited enhanced extravasation through the BM compared to unmodified GD2.CAR T cells (5.90e6 um^2^ ± 5.66e5 um^2^). Further, we observed increased overall infiltration of MMP7-GD2.CAR (1.81e6 um^2^ ± 3.57e5 um^2^, day 4, *p < .01*) and OPN-GD2.CAR T cells (1.12e6 um^2^ ± 2.24e5 um^2^, *p < .05*) into tumor droplets compared to unmodified GD2.CAR T cells (6.70e5 um^2^ ± 2.69e5 um^2^) (**Fig 3B, 3E**). Tumor droplet infiltration was verified by confocal microscopy (**Supplementary** Fig 2A).

**Figure 3.**
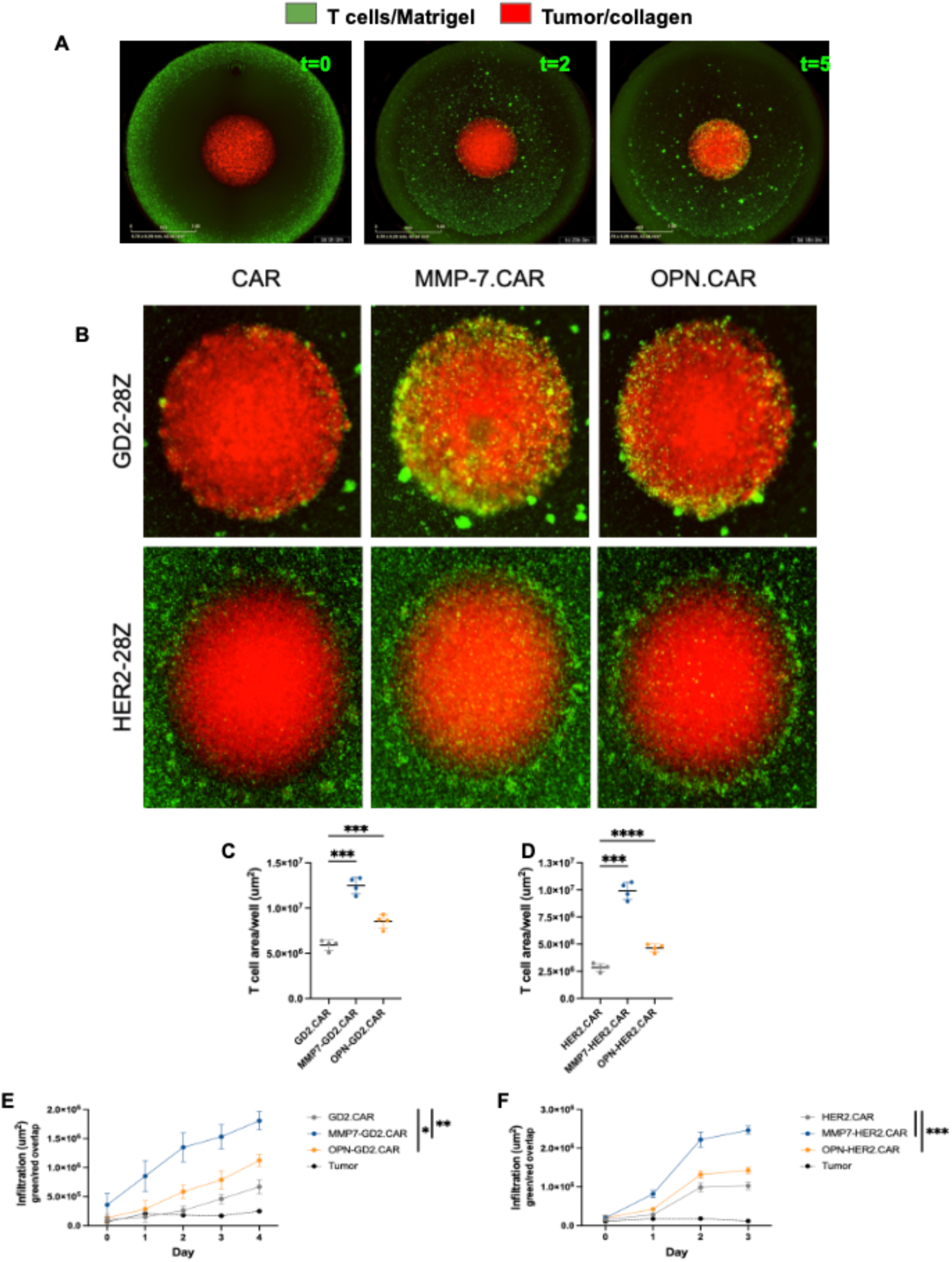
MMP-7 and OPN overexpression increase CAR T infiltration *in vitro*. **A.** Sample images of the Halo infiltration assay on day 0, day 2, and day 5. GFP-expressing CAR T cells were resuspended in Matrigel and plated in a halo on the perimeter of a well, while tumor target cells were stained red, resuspended in collagen type I, and plated as a droplet in the center of the well (t=0). CAR T cells extravasate out of the Matrigel (t=2), migrate across the well, and then infiltrate the tumor droplet (t=5). **B.** Images of tumor droplets infiltrated by unmodified GD2.CAR, MMP7- GD2.CAR or OPN.GD2.CAR T cells (top) on day 5 of the Halo assay, or unmodified HER2.CAR, MMP7-HER2.CAR or OPN.HER2.CAR T cells (bottom) on day 3 of the Halo assay. **C-D.** Extravasation of modified (C) GD2.CAR T cells at t=12h, and (D) HER2.CAR T cells at t = 6h measured as T cell area/well (mean, ± SD, *n = 4*, one-way ANOVA with Dunnett’s correction). **E-F.** Infiltration of modified (E) GD2.CAR T cells over four days and (F) HER2.CAR T cells over three days, measured as the area where CAR T cells (green) overlap tumor droplets (red) (mean, SEM, *n = 4*, mixed-effects analysis with multiple comparisons). *\*p < .05, **p < .01, ***p < .001, ****p < .0001*.

We next asked if MMP-7 or OPN overexpression could improve the infiltration of CAR T cells targeting other solid-tumor antigens. We therefore overexpressed MMP-7 or OPN in T cells expressing a HER2-41BBZ CAR (HER2.CAR) and used our Halo assay to compare the extravasation and infiltration of MMP7-HER2.CAR, OPN-HER2.CAR and unmodified HER2.CAR T cells. We observed the same pattern of enhanced extravasation of MMP7-HER2.CAR (9.92e6 um^2^ ± 7.91e5 um^2^, t = 6h, *n = 4, p < .001*) and OPN-HER2.CAR T cells (4.64e6 um^2^ ± 3.62e5 um^2^, *p < .0001*) compared to unmodified HER2.CAR T cells (2.85e6 um^2^ ± 3.65e5 um^2^) (**Fig 3D**). Total tumor droplet infiltration was also increased in MMP7-HER2.CAR (2.46e6 um^2^ ± 2.15e5 um^2^, day 3, *p <.001*) and OPN-HER2.CAR T cells (1.43e6 um^2^ ± 1.61e5 um^2^, *p < .001*) compared to unmodified HER2.CAR T cells (1.03e6 x um^2^ ± 1.97e5 um^2^) (**Fig 3B, 3F**). While we found that the expression of MMP-7 or OPN enhanced CAR T cell infiltration, we did not observe any differences in tumor invasion in the presence of these secreted proteins (**Supplementary** Fig 3A).

Having confirmed that MMP-7 and OPN enhanced tumor infiltration by GD2.CAR T cells *in vitro*, we characterized the CD8+ fraction of these cells using gene expression analysis. While we did not observe differences in genes associated with activation or exhaustion (**Supplementary** Fig 4A-B), surprisingly, there was a broad upregulation of MMPs in MMP7-GD2.CARTs, and to a much lesser extent, OPN-GD2.CAR T cells compared to unmodified GD2.CAR T cells (**Fig 4A**). To confirm these differences at the protein level, we stimulated MMP7.GD2-CAR, OPN.GD2-CAR, unmodified GD2.CAR T and GFP-transduced T cells with LAN-1 tumor cells and measured the secretion of select MMPs. We found that the secretion of MMP-7, MMP-1, MMP-3 and MMP-8 indeed was higher in MMP7-GD2.CAR (*n = 4, p < .01*) than unmodified GD2.CAR T cells, while only MMP-3 secretion was increased in OPN-GD2.CAR T cells (*p < .05*) (**Fig 4B**).

**Figure 4.**
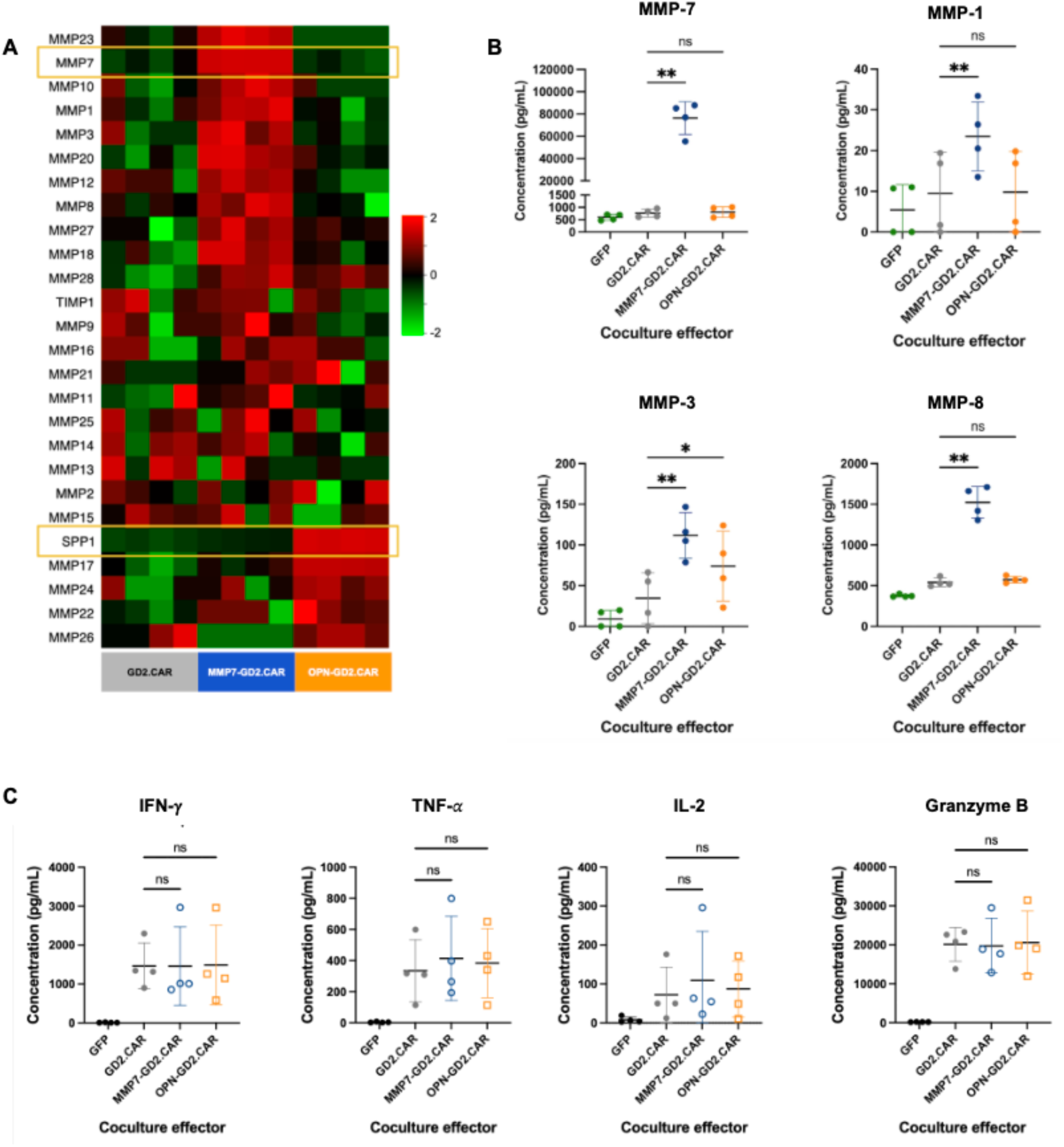
MMP7-GD2.CAR and OPN-GD2.CAR T cells broadly upregulate MMPs. **A.** Heatmap of *MMP* and *SPP1* expression in CD8+ MMP7-GD2.CAR, OPN-GD2.CAR and GD2.CAR T cells at rest (day 9 post-activation, *n = 4*). **B.** Secretion of select MMPs by tumor-stimulated MMP7-GD2.CAR, OPN-GD2.CAR and unmodified GD2.CAR T cells. **C.** Secretion of cytokines by tumor-stimulated MMP7-GD2.CAR, OPN-GD2.CAR and unmodified GD2.CAR T cells. Data presented is the mean, SD, *n = 4*, one-way ANOVA with Dunnett’s correction. *\*p < .05, **p < .01*.

In contrast to the expression of MMPs, neither MMP-7 nor OPN had any effect on the secretion of IFN-𝛾, TNF-𝛼, IL-2 or granzyme B after tumor-stimulation of GD2.CAR T cells, suggesting that GD2.CAR T cells maintained stable expression of key immune cytokines when MMP-7 or OPN was coexpressed (**Fig 4C**).

*OPN-GD2.CAR T cells exhibit superior infiltration and tumor clearance in a xenograft NB model.* To determine how MMP-7 or OPN affected *in vivo* tumor infiltration, we used a xenograft model of NB to evaluate tumor clearance and survival after GD2.CAR T infusion. We injected firefly luciferase transduced LAN-1 (LAN-1-FFLuc) cells into NSG mice, and allowed tumors to grow until they were 1.0 cm in diameter, then gave a sub-curative dose of GD2.CAR T cells by tail vein injection. We selected subcutaneous models because ECM that is histologically equivalent to patient disease is not achieved until tumors are approximately 1 cm in diameter^32^, a size that cannot be achieved in kidney-implanted orthotopic models due to limitations imposed by animal comfort. When we tracked tumor burden by IVIS imaging, we found that GD2.CAR or MMP7- GD2.CAR T cell treated mice had an initial reduction in tumor burden between days four and seven post-treatment, followed by the gradual outgrowth of tumors. In contrast, tumor burden in mice treated with OPN-GD2.CAR T cells showed a sharp decline on day four post-treatment that was sustained over time (**Fig 5B**). On day 29 post-treatment, tumor burden was 94% lower in the OPN-GD2.CAR T treated mice (6.57e7 p/s ± 1.53e8 p/s, mean, SD, *n = 12, p < .05*) compared to GD2.CAR T (1.06e9 p/s ± 1.63e9 p/s, *n = 9*) treated mice, however, we did not observe reduced tumor burden in MMP7-GD2.CAR T treated mice (**Fig 5C**). Even though the dose of CAR T cells was sub-curative, reduced tumor burden in OPN-GD2.CAR T cell treated mice resulted in an increase in overall survival by a median of 148 days (*n = 9-12, p < .05*) (**Fig 5D**).

**Figure 5.**
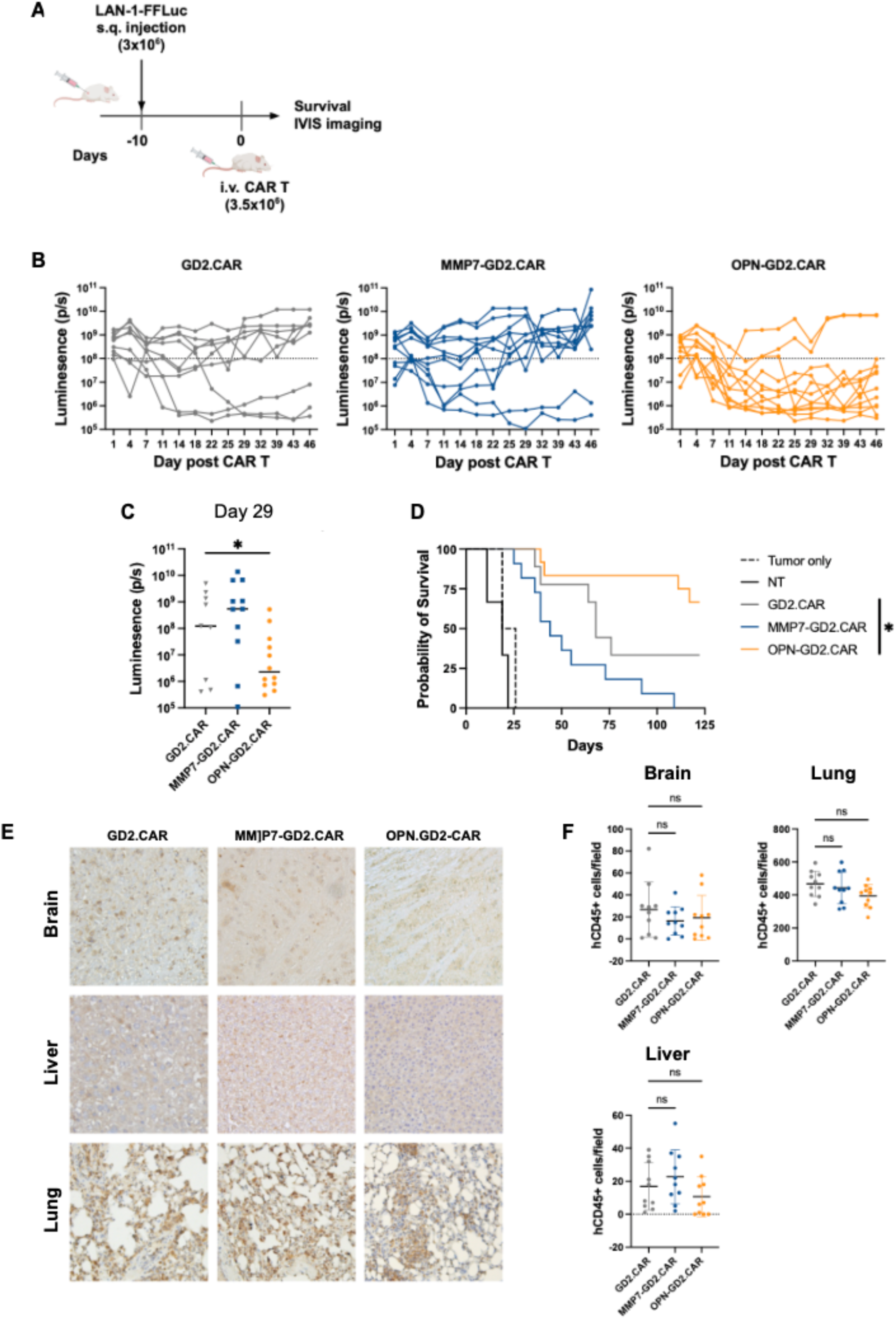
OPN-GD2.CAR T cells exhibit superior tumor control in a xenograft model of NB. **A.** Schematic of xenograft model of NB and CAR T cell dosing. Firefly luciferase transduced LAN-1 (LAN-1-FFLuc) cells were injected subcutaneously into the flanks of NSG mice. When tumors reached 0.75-1 cm in diameter, 3.5e6 GD2.CAR, MMP7-GD2.CAR or OPN-GD2.CAR T cells were infused by tail vein injection. **B-C.** Plots of tumor burden over 46 days (**B**), and on day 29 post CAR T infusion (**C**) (median, unpaired t test, *n = 9-12*). **D.** Kaplan-Meier survival curve for days 0- 125, with significance determined by log-rank test (Mantel-Cox). Results shown are from two separate experiments and donors. **E.** IHC of brain, lung, and liver tissue sections stained for human CD45, collected from mice treated with GD2.CAR, MMP-7.GD2.CAR or OPN-GD2.CAR T cells (*n = 5*). **F.** Quantification of CD45+ cells in stained brain, lung, and liver tissue (mean, SD, *n = 10*, one-way ANOVA with Dunnett’s correction). *\*p < .05, **p < .01*.

Because MMP7- and OPN-GD2.CAR T cells exhibited enhanced infiltration *in vitro*, we wanted to ensure that our modified cells did not infiltrate healthy tissues. We therefore collected brain, lung, and liver tissue from mice sacrificed at the humane experimental endpoint of our study and stained sections of these tissues for human CD45 (**Fig 5E**). T cells were then quantified by counting. We observed no significant differences in CAR T cell infiltration into healthy tissues when comparing the number of infiltrating MMP7-GD2.CAR, OPN-GD2.CAR and unmodified GD2.CAR T cells (**Fig 5F**). These data suggest that the expression of MMP-7 or OPN in CAR T cells does not augment non-specific infiltration.

Once we established that forced OPN expression induced the greatest tumor control by GD2.CAR T cells, we asked whether that effect was indeed associated with improved tumor infiltration. To evaluate this, we modified our existing mouse model to focus on T cell localization (**Fig 6A**). Here, we used firefly luciferase (FFLuc)-transduced GD2.CAR (GD2.CAR-FFLuc), MMP7-GD2.CAR (MMP7-GD2.CAR-FFLuc) or OPN-GD2.CAR (OPN-GD2.CAR-FFLuc) T cells to treat established WT LAN-1 tumors (**Fig 6A**). Each group of mice was then split into two cohorts. The first cohort was imaged every two to three days to track T cell localization by bioluminescence. The second cohort was sacrificed five days post-CAR T treatment, and tumors were collected from sacrificed mice, sectioned and stained for collagen type I and hCD3. Intratumoral CAR T cells were then quantified by counting CD3+ cells within the dense collagenous margins of the tumor.

**Figure 6.**
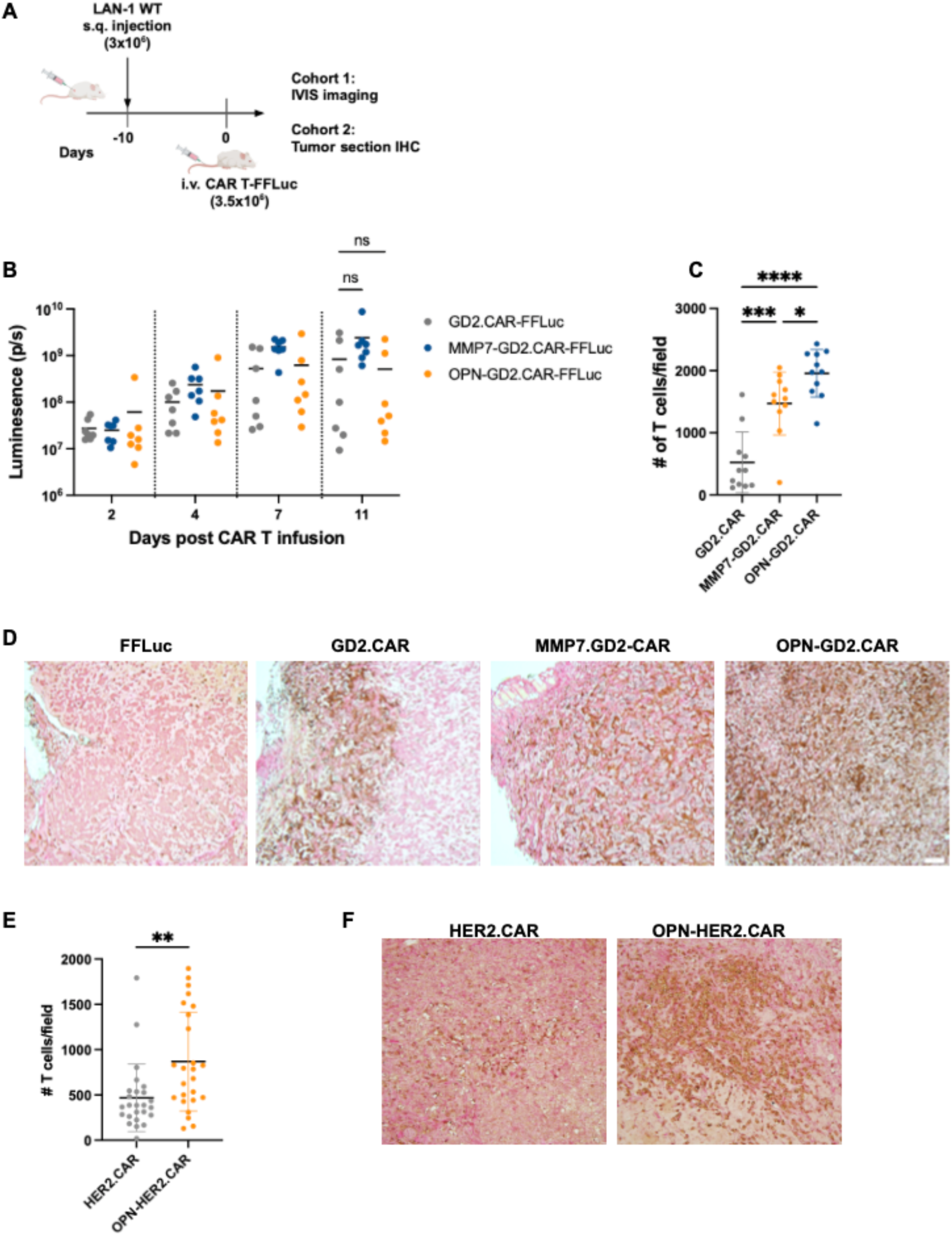
OPN overexpression in CAR T cells enhances infiltration into solid tumors. **A.** Schematic of mouse model for CAR T localization. NSG mice were subcutaneously injected with 3e6 WT LAN-1 cells and tumors were allowed to grow until they reached 0.75-1.0 cm in diameter. Mice were then treated with 3.5e6 GD2.CAR-FFLuc, MMP7-GD2.CAR-FFLuc or OPN-GD2.CAR- FFLuc T cells by tail vein injection, and each group of mice was split into two cohorts. In one cohort, IVIS imaging was used to track T cell localization over 11 days post-treatment. In the second cohort, five days post-treatment, tumors were collected and stained for collagen type I and hCD3. **B.** T cell luminescence at the tumor site (mean, *n =* 7, two-way ANOVA with Dunnett’s correction). **C-D.** IHC of NB tumors stained for collagen type I (red) and hCD3 (brown). Random fields within the tumor margin were imaged and the number of T cells per field was counted. **C.** Quantification of infiltrating CAR T cells in NB tumors (mean, SD, *n = 11*, one-way ANOVA with Tukey’s correction). **D.** Representative sections of treated NB tumors. **E-F.** Xenograft BC model in which 5e6 MDA.MB.453 breast tumor cells were injected sq into NSG mice. Tumors were allowed to develop until they reached at least 1.0 cm in diameter. Mice were then treated with 3.5e6 HER2.CAR or OPN-HER2.CAR T cells. Five days post-treatment, mice were sacrificed and tumors were collected, sectioned and stained as in our NB model. **E.** Quantification of intratumoral OPN- HER2.CAR and HER2.CAR T cells (mean, SD, *n =25*, unpaired t test). *\* p < .05, **p < .01, ***p < .001, **** p < .0001*.

We observed comparable levels of bioluminescence at the tumor site across all three conditions, suggesting equivalent trafficking of CAR T cells to the tumor site on day 11 (**Fig 6B**). However, immunohistochemistry (IHC) staining revealed a drastic increase in intratumoral CAR T cells in OPN-GD2.CAR (1,957 cells/field +/- 381.4 cells, *n = 11*) and MMP7-GD2.CAR (1,470 cells/field +/- 507.1 cells, *n = 11*) T cell treated mice compared to mice treated with unmodified GD2.CAR T cells (522.8 cells/field +/- 490.1 cells, *n = 11*) (**Fig 6C**). Further, there were notable differences in the pattern of distribution of CAR T cells within the tumors between conditions. GD2.CAR T cells infiltrated from the perimeter of the tumor inward, with the majority of cells being localized near the tumor margins, associated with collagen-dense regions (**Fig 6D**). In contrast, MMP7-GD2.CAR and OPN-GD2.CAR T cells showed an even distribution throughout the tumor, penetrating directly into and throughout the tumor core (**Fig 6D**).

Compared to unmodified GD2.CAR T cells, MMP7-GD2.CAR T cells exhibited equivalent expansion, cytotoxicity, phenotypic ratios and effector molecule release upon stimulation *in vitro*, but failed to clear tumors *in vivo*. We hypothesized that the lack of efficacy of MMP7- GD2.CAR T cells may be due to exhaustion. Therefore, we looked for the presence of PD-1, LAG- 3, TIM-3 and CD39 on TILs isolated from NB tumors collected from treated mice, but saw no significant differences between MMP7-GD2.CAR and unmodified GD2.CAR T cells across these markers (**Supplementary** Fig 5A).

### Overexpression of OPN may be broadly applicable to other CARs and tumor types

Because OPN overexpression enhanced both GD2.CAR and HER2.CAR T cell infiltration *in vitro*, we investigated whether OPN-HER2.CAR T cells would exhibit enhanced infiltration *in vivo*. To examine OPN-HER2.CAR T infiltration, we used a xenograft model of BC. We established subcutaneous MDA-MB-453 tumors in NSG mice and infused HER2.CAR, OPN-HER2.CAR or NT T cells by tail vein injection on day 14. Five days post-infusion we collected the tumors and quantified CAR T cell infiltration using IHC and counting. Our data showed that OPN-HER2.CAR T cells (868.6 cells +/- 546.6 cells, mean, SD, *n = 25*) exhibited enhanced infiltration into BC tumors compared to unmodified HER2.CAR T cells (469.5 cells +/- 372.1 cells, *n = 25*) (**Fig 6E**). Furthermore, the pattern of infiltration was similar to NB tumors, where unmodified HER2.CAR T cells showed lower infiltration into the center of the tumor compared with OPN-HER2.CAR T cells (**Fig 6F**), suggesting the expression of OPN may improve the infiltration of other solid-tumor targeting CARs.

## DISCUSSION

Our data demonstrate that GD2.CAR T cells face infiltration restrictions due to the dense, tumor-specific ECM associated with bulky tumors. Forced expression of OPN in these cells enhanced their infiltration, resulting in improved tumor clearance and survival. These findings highlight a novel approach to overcome a key barrier to CAR T cell efficacy in solid tumors.

Solid tumors comprise 93%^37^ of all cancer deaths, but T cell therapies for solid tumors universally face challenges posed by the tumor ECM. Patient-derived samples reveal that endogenous T cells aggregate within collagen-dense regions, mirroring findings from our transwell invasion assays. This study is the first to show that tonically-signaling GD2.CAR T cells, which are widely used for solid tumor treatment, are even more restricted in ECM traversal. These characteristics make tonically-signaling GD2.CAR T cells prime candidates for optimization through gene modification.

OPN stands out as a promising gene modification target due to its dual role in cell movement. T cells utilize two primary modes of movement to infiltrate tumors: extravasation, where they exit blood vessels into tumor tissue, and interstitial movement, which involves navigating the ECM to reach tumor cells. Both processes rely on forces generated by the cortical actin cytoskeleton, which is anchored to the ECM through interactions with integrins or adhesion molecules^38,39^. As a secreted protein, OPN binds to ECM fibers such as collagen (types I-V) and fibronectin. There it acts as a ligand for integrins, facilitating interstitial movement, and for CD44, which is crucial for T cell extravasation into inflamed tissues. Intracellularly, a distinct isoform of OPN, translated from an alternate start site, interacts with ERM proteins which link the cortical actin cytoskeleton to the cytoplasmic tails of integrins and CD44^40,41^. These connections enable the cytoskeleton to exert the forces necessary for translocation during both extravasation and interstitial movement. Thus, OPN serves as a vital anchoring point, enhancing T cell motility and tumor infiltration from both inside and outside the cell. By leveraging OPN’s ability to enhance both extravasation and interstitial movement, we achieved significant improvements in GD2.CAR T cell infiltration.

OPN has been associated with metastatic cancer spread^42,43^, but our results show no evidence of increased non-specific infiltration into healthy tissues such as the brain, lungs, or liver when using MMP-7 or OPN-modified GD2.CAR T cells. Additionally, concerns about OPN’s role in promoting tumor invasion were mitigated by findings that CAR T cell-secreted OPN did not increase tumor invasiveness in our *in vitro* models. Unlike tumor-derived soluble OPN, CAR T cell-derived OPN is ECM-associated, with its function heavily influenced by post-translational modifications^44^. Further, no tumor metastasis was observed in our xenograft models treated with OPN-expressing CAR T cells, supporting the safety of this approach.

The efficacy of GD2.CAR T cell therapy has been limited in clinical cases of bulky disease, despite promising preclinical results. This discrepancy may stem from the limitations of *in vivo* models in studying tumor infiltration. While orthotopic models mimic the biological niche of clinical tumors, in mice they often fail to achieve the tumor size and ECM density seen in bulky disease^45^. Our subcutaneous models, chosen for their ability to develop collagen-rich tumors of sufficient size, better replicate the infiltration challenges faced in clinical settings. For future studies, orthotopic canine or non-human primate models may offer even greater biological relevance for infiltration studies.

Interestingly, MMP7-GD2.CAR T cells exhibited enhanced infiltration *in vivo* but failed to clear tumors. This paradox may be explained by the broad upregulation of matrix metalloproteinases (MMPs) we observed, which could lead to the cleavage of critical components of the immune synapse, such as LFA-1 and ICAM^46,47^. Disruption of the immune synapse may not affect *in vitro* cytotoxicity assays, where effector-to-target ratios favor T cells, but it could impede efficacy in the tumor microenvironment.

GD2.CAR T cell therapy has shown promise in treating NB but remains ineffective against bulky disease. While CAR T cell trafficking has been widely investigated, infiltration remains poorly understood. Our findings underscore the importance of studying the tumor-specific ECM as a barrier to T cell movement. Enhanced infiltration, as achieved through OPN expression, could alleviate two major contributors to T cell therapy failure, low persistence and exhaustion: By facilitating deeper tumor penetration by a broader population of CAR T cells, OPN expression reduces the serial killing burden on individual T cells that contributes to exhaustion and poor persistence. This study lays the groundwork for future investigations into gene modifications aimed at enhancing CAR T cell infiltration and effectiveness in solid tumors, highlighting forced expression of OPN as a promising approach to optimize GD2.CAR T cell therapy for bulky tumors.

## MATERIALS AND METHODS

### Cell lines

Peripheral blood was obtained from healthy donors under a Baylor College of Medicine (BCM) Institutional Review Board-approved protocol. Donors provided informed written consent, and research was guided by the Declaration of Helsinki, the International Ethical Guidelines for Biomedical Research Involving Human Subjects (CIOMS), Belmont Report, and Common Rule. Peripheral blood mononuclear cells (PBMCs) were then isolated by density gradient centrifugation using Lymphoprep solution (Stemcell Technologies, #07801). In experiments using freshly-isolated T cells (FI-T), T cells were isolated from PBMCs by negative selection using Miltenyi Biotec’s Pan T isolation kit (#130-096-535) per the manufacturer’s instructions and used immediately. For all other experiments, PBMCs were frozen and stored.

293T cells were purchased from ATCC (#CRL-3216), and LAN-1 cells were purchased from Sigma Aldrich (#06041201-1VL). LAN-1 cells were retrovirally transduced to express GFP and firefly luciferase to generate LAN-1-GFP-FFLuc. MDA.MB.453 cells were a generous gift from Valentina Hoyos. All cell lines were maintained in HyClone Dulbecco’s Modified Eagle Medium (Cytiva, #SH30081.01) supplemented with 10% FBS and 2 mmol/L GlutaMAX (Gibco, #35050-061). All cell lines in culture were checked for mycoplasma using an enzyme-based assay (Lonza), every eight weeks and before freezing.

### Retroviral vectors and virus production

To generate the MMP-7 and MMP-14 vectors, sequences encoding MMP-7 or MMP-14, followed by T2A and mEmerald were cloned in-frame into an SFG retroviral vector with In-Fusion cloning (Takara Bio In-Fusion HD Cloning Kit, #639649). To generate OPN vectors, sequences encoding OPN (isoform B) followed by T2A and either mEmerald or truncated CD19 (tCD19) were cloned in-frame into an SFG vector. Retroviral vectors encoding HER2-28Z^26^, GD2-41BBZ^27^, GD2-28Z^27^, CD19-41BBZ^28^, and CD19-28Z^28^ CARs have been described previously described. The GFP plasmid MSCV-IRES-GFP was purchased from AddGene (#20672DSHO).

Lipofectamine 3000 (Thermo Fisher Scientific #L3000001) was used according to the manufacturer’s instructions to transfect 293T cells growing in complete DMEM with the SFG plasmid encoding our gene of interest including LTRs and packaging signals, a PegPAM plasmid encoding MoMLV gag-pol, and an RDF plasmid encoding RD114 envelope. Medium was replaced 24 hours after transfection. Viral supernatant was harvested and filtered with a 0.45-um filter (PALL Life Sciences, #4654) 48- and 72 hours post-transfection. Virus collected at 48 hours was stored at 4^1^C until the 72-hour virus was harvested, then these supernatants were pooled and used immediately for transduction.

### Generation of ATCs

Non-tissue culture treated 24-well plates (Falcon, #351147) were coated with 0.5 ug per well of CD3 antibody produced in-house by the OKT3 hybridoma (ATCC, #CRL-8011) and CD28 antibody (BD Pharminogen, #555725, clone CD28.2), both resuspended at 1 ug/mL in sterile water. Plates were incubated overnight at 4°C and the antibody mixture was aspirated before PBMCs were plated. CTL medium, a 1.5:1 mixture of RPMI 1640 (Hyclone, #SH3002701) and Click’s medium (Fujifilm Irvine Scientific, #92705) supplemented with 10% FBS and 2 mmol/L GlutaMAX (Thermo Fisher Scientific, #35050061) was used to culture ATCs and T cell lines. PBMCs were thawed, resuspended in CTL medium supplemented with 10 ng/mL IL-7 and IL-15 (R&D Systems, #204IL), and added to coated plates at 1e6 cells per well. ATCs were split and cytokines were replenished every 2-3 days. Expansion was tracked by counting cells using trypan blue exclusion on a TC10 cell counter (BioRad).

### Retroviral transduction

Non-tissue culture treated 24-well plates were coated with 3.5 ug of RetroNectin (Takara, #T100B) in 0.5 mL of Dulbecco’s PBS (Sigma-Aldrich, #D8537) per well and plates were incubated at 4^1^C overnight. The next day, the RetroNectin solution was removed and replaced with 2 ml per well of fresh retroviral supernatant, or for nontransduced cells complete DMEM. Plates were centrifuged at 1,000g for one hour at room temperature, the viral supernatant was aspirated and ATCs were plated at 2.5e5 cells per well in CTL medium supplemented with 10 ng/mL IL-7 and IL-15. For MMP7-GD2, OPN-GD2 and GFP-GD2, T cells were transduced sequentially with the GD2- 28Z CAR on day 2 post-activation, and with MMP7.T2A.GFP, OPN.T2A.GFP, OPN.T2A.tCD19 or GFP on day 3 post-activation. All other conditions (GD2-41BBZ, GD2-28Z, GFP, delta CD19, CD19- 41BBZ, CD19-28Z) were transduced on day 2 post-activation. Two days after transduction, transduction efficiency was evaluated by flow cytometry and T cells were split and moved to tissue-culture treated plates.

### Flow cytometry

Transduction was measured by flow cytometry using untransduced or GFP transduced cells as controls. Antibodies were added to 5e5 cells and incubated for 15 minutes in the dark at room temperature. Cells were washed twice with PBS, and analyzed on a BD FACS Canto cytometer. Analysis was done using FlowJo. GD2-CAR expression was measured using the idiotype antibody 1A7 conjugated to AlexaFluor-647. Transduction of MMP-7 was measured by GFP expression, and transduction of OPN was measured by either GFP expression or using a PE-conjugated antibody to CD19 (BioLegend, #302208, clone HIB19). HER2-CAR expression was detected using an AF647-conjugated antibody against mouse IgG_1_ F(ab’)_2_ (Jackson Immuno Research Laboratory, #315-605-006). Transduction of the CD19-CARs (CD19-41BBZ, CD19-28Z, and ΔCD19) was detected using a PE-conjugated antibody against CD271 (NGFR) (BioLegend, #345105, clone ME20.4). The proportions of CD4+ and CD8+ T cells was determined using CD3-APC (BD Pharmingen, #555342), CD4-PerCP (BioLegend, #317431, cone OKT4), and CD8-FITC (BioLegend, #301050, clone RPA-T8). Cell phenotype was determined using CCR7-PE (BioLegend, #353204, clone G043H7), CD45RO-PE-Cy7 (BioLegend, #304230, clone UCHL1) and defined as effector memory (T_EM_ CD45RO+/CCR7-), central memory (T_CM_ CD45RO+/CCR7+), effector (T_EFF_ CD45RO-/CCR7-) or naïve (T_N_ CD45RO-/CCR7+) T cells. Exhaustion markers were detected using CD279- BV650 (BioLegend, #329950, clone EH12.2H7), CD366-PE (BioLegend, #345006, clone F38-2E2), CD223-APC-Cy7 (BioLegend, #369347, clone 11C3C65), and CD39-BV421 (BioLegend, #328214, clone A1).

### Immunohistochemistry

Tumors and organs were fixed in 10% neutral buffered formalin (Globe Scientific, #6527FL) for at least 48 hours, then embedded in paraffin and sectioned into 4um sections. Staining was done by the Texas Children’s Hospital Pathology department. Brain, lung, and liver sections were stained for human CD45 using a primary mouse anti-human CD45 monoclonal antibody (Dako, #M070101-2, clone 2B11 + PD7/26) followed by a goat anti-mouse secondary (Vector Laboratories, #MP-7452). Tumor sections were stained for human CD3 (Dako, #A045201-2) on the BOND III (Leica, #21.2201) automated staining system using protocol “F” with BOND Polymer Refine Detection (Leica, #DS9800), followed by collagen staining with picrosirius red (Polysciences, Inc., #24901-250) according to manufacturer’s instructions. Tumor infiltration by CAR T cells and non-specific infiltration into organs were visualized at 20X on a Leica DMi8 microscope using CellSense software. T cells were quantified in 12 random fields in tumors and 24 random fields in organs by counting using ImageJ (Version 1.53t).

### Transwell invasion assay

Matrigel (Corning, #356234) was diluted to the indicated concentrations with pure RPMI, or for assays comparing the infiltration of FI-T and CAR T cells, diluted 1:10. T cells were resuspended at 2.5e6 cells/mL in diluted Matrigel, and 100 uL was plated in Thincert transwell inserts (8 um pore size, Greiner #662638) and placed inside non-tissue culture treated 24 well plates (Falcon, #351147). Plates were incubated at 37^1^C for 60 minutes to allow the Matrigel to polymerize. Coated wells were then topped with 50 uL of pure RPMI, and 750 uL of complete CTL medium was added to the bottom of the wells. Conditions were plated in quadruplicate, then incubated overnight at 37^1^C. The next day, the matrix was dissociated by incubation with collagenase [2mg/mL] for 20 minutes at 37^1^C. The non-infiltrating cells isolated from the dissociated matrix and the infiltrating cells collected from the bottom of each well were counted. The % invasion was calculated as: (number of invading cells/(number of invading + non-invading cells)) x 100%. For tumor invasion assays the procedure was the same except Matrigel was diluted, with pure DMEM, and polymerized gels were covered with pure DMEM. The bottom chambers of the transwells were filled with 750 uL of supernatants collected from cocultures of GFP, GD2.CAR, MMP7-GD2.CAR or OPN.GD2.CAR T cells and LAN-1 tumor cells, that had been plated at a 2:1 ratio in complete CTL medium for 24 hours.

### Gene expression analysis of published datasets

Gene expression datasets GSE117230, GSE77043, GSE183349, and GSE8476 were accessed through the National Center for Biotechnology Information’s (NCBI) Gene Expression Omnibus (GEO) database. Analysis was performed using GEO2R, which uses GEOquery and limma to perform differential expression analysis. Infiltrating populations were defined as the test groups, while non-infiltrating populations served as control groups. The top 250 differentially expressed genes in infiltrating populations were then ranked by log_2_ fold change; genes for non-coding RNAs were excluded. The top 50 differentially expressed genes from each study were then cross-referenced to identify genes that were upregulated in more than one study. Genes involved in cell movement were identified from the list of commonly expressed genes in infiltrating populations.

### Cytotoxicity

Chromium release cytotoxicity assays were performed using LAN-1 neuroblastoma cells as targets. To label targets, 1e6 LAN-1 cells were pelleted and resuspended in 200 uL of complete DMEM containing ^51^Cr, and incubated at 37°C for one hour. The cells were then washed three times and resuspended in 10 mL of CTL medium. Effector T cells included nontransduced ATCs, and GD2-28Z, MMP7.GD2-28Z, and OPN.GD2-28Z CAR Ts. Effector and target cells were then cocultured in a 96-well plate for six hours at various ratios (40:1, 20:1, 10:1, 5:1), spun down at 400g for three minutes, and 100 uL of supernatant was used for analysis. Target cells plated without effectors served as the control for spontaneous release, and targets lysed using 0.1% Triton X were used as the control for maximum release. Samples were analyzed for the amount of chromium released (counts per minute, CPM) using a gamma counter (PerkinElmer 2470 WIZARD2). Specific lysis was calculated as (sample release – spontaneous release)/(maximum release – spontaneous release). All conditions were plated in triplicate.

### ELISAs

To measure secreted protein levels of MMP-7 and OPN, nontransduced T cells, GD2-28Z, MMP7- GD2 and OPN-GD2 CAR T cells generated from 3 healthy donors were each plated at 1e6 cells per well in CTL medium supplemented with 10 ng/mL IL-7 and IL-15 and incubated at 37^1^C for 24 hours. The next day, supernatants were collected, centrifuged at 800g for 10 minutes, and stored at -80^1^C. All conditions were plated in triplicate ten days after activation. Samples were then thawed at room temperature, and quantified for MMP-7 (BioVendor, #QZBMMP7H) or OPN (R&D Systems, #DOST00) by ELISA according to manufacturer’s instructions.

### Halo infiltration assay

#### Collagen preparation

A 2.5 mg/mL solution of collagen I was prepared from collagen I (rat tail, Corning, #354236), 10X DPBS (Sigma, #D1408-500ML), 1N NaOH (Sigma, #S2770-100ML), and cRPMI (RPMI supplemented with 2mmol/L GlutaMAX (Gibco, #35050-061) and 10% FBS) at the following volumes of chilled reagents, with NaOH being added last, and the final solution being kept on ice until needed:

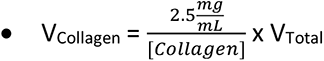

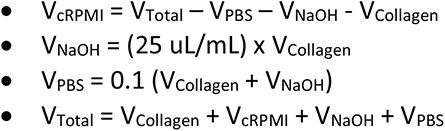

#### Cell staining and plating

LAN-1 tumor cells were stained with CellBrite Red (Biotium, #30023) according to manufacturer’s directions, and resuspended at 1e5 cells/uL in the 2.5 mg/mL collagen I solution, and a 1 uL tumor droplet was plated in the center of the well in 96-well tissue culture treated flat bottom plates (Corning, #3585). T cells transduced with GFP were then resuspended at 40e6 cells/mL in a 4:3 mixture of Matrigel (Corning, #356234) and cold cRPMI, and 5 uL of T cells was plated on the perimeter of each well containing tumor droplets. Matrices were polymerized by incubation at 37^1^C for 15 minutes, then covered with 150 uL cRPMI and placed in an Incucyte S3 Live-Cell Analysis Instrument (Sartorius, Goettingen, Germany) for imaging. A 4X objective was used to take whole well images every 6 hours in the red and green channels. For image capture, spectral unmixing was used and set as %R contributes to G is 10%.

#### Image analysis

To analyze the green channel, top hat segmentation was used with a radius of 100 um, the threshold was set to 2.5 GCU, and the edge split option was selected with an edge sensitivity of -10. No hole fill was used for green channel images. To analyze the red channel, top hat segmentation was used with a radius of 100 um, the threshold was set to 2.0 GCU, and the edge split option was selected with an edge sensitivity of -40. The hole fill setting in the red channel was set to 8,000 um^2^ to create a solid area where T cell overlap could be detected. To measure infiltration, a mask was created using the noted settings to identify T cells in the green channel, and a mask was created to identify the tumor droplet in the red channel. Infiltration was measured as the area (um^2^) where the T cell mask overlapped the tumor droplet mask. All conditions were plated in quadruplicate. Extravasation was measured as the total area (um^2^) of T cells in each well at six hours after plating.

### Confocal imaging of Halo assay

96-well flat-bottom plates (Corning Costar, #3595) were imaged live once a day at (37C/ 5% CO2) for 5 consecutive days using a Yokogawa CV8000 spinning disk confocal microscope. Images were acquired using both a 4X/0.16NA and 20xW/1.0NA objectives. The 4X images covered the whole well with four fields of 50 um optical sections (200um Z-stack). With the 20X water immersion lens, nine fields of view were acquired (sixteen 2.5 um optical sections). Images were captured in both fluorescence and brightfield modes, using the following laser/filter combinations: 488 nm/525 nm (Ex/Em) and 640 nm/67 6nm (Ex/Em). Laser power and exposure times were adjusted to capture sufficient signal of at least 10/1 signal/noise ratio due to the fact that T cells used for different experimental conditions showed varying fluorescent intensities in the 488 nm channel. 4X images were max intensity-projected using CellPathfinder software (Yokogawa). In the 20X image datasets, the middle z plane was chosen to represent T cell infiltration within the tumor droplet. Histograms were adjusted in ImageJ/Fiji for all TIFF images to show comparable brightness from each channel.

### Luminex

2.5e5 GFP, GD2.CAR, MMP7-GD2.CAR or OPN-GD2.CAR T cells (day 9 post-activation) were plated at a 2:1 ratio with LAN-1 cells for 24 hours, and supernatants were collected, centrifuged, and stored as aliquots at -20^1^C. Aliquots were thawed quickly and analytes were measured using a Human Luminex Discovery Assay (BioTechne, #LXSAHM-19) according to the manufacturer’s instructions. Plates were read on a Luminex 200 instrument.

### Xenograft mouse models

All animal experiments were conducted under a BCM Institutional Animal Care and Use Committee approved protocol. In our NB survival model, 6-week old female NOD- scidIL2Rgamma^null^ (NSG, The Jackson Laboratory, #005557) mice were given subcutaneous injections of 3e6 LAN1-FFLuc cells, resuspended in 100uL of Matrigel, into their right flanks. Tumor growth was monitored by caliper measurement. When tumors reached 0.7-1.0 cm in diameter, mice were distributed by size into treatment groups to minimize differences in mean tumor diameter. Mice were then treated with 3.5e6 nontransduced T cells or GFP, GD2.CAR, MMP7-GD2.CAR, OPN-GD2.GD2.CAR T cells by tail vein injection. In T cell localization experiments, tumors were established with WT LAN-1 cells and mice were treated with FFLuc, GD2.CAR-FFLuc, MMP7-GD2.CAR-FFLuc, or OPN-GD2.CAR-FFLuc T cells. The xenograft model of BC followed the same protocol, except tumors were established with 5e6 MDA.MB.453 cells.

T cells used for these experiments were generated, frozen on day 5 after activation, then thawed, expanded for four days in 10 ng/mL of IL-7 and IL-15, and infused. Tumor burden was measured twice a week with calipers and bioluminescent imaging (BLI, Xenogen, IVIS, Small Animal Core Facility, Texas Children’s Hospital). For BLI, mice were anaesthetized with isoflurane (Covetrus, #029405), injected intraperitoneally with D-luciferin (Perkin Elmer, #122799) and imaged after 10 minutes. Mice were euthanized when tumors reached 1.5 cm in diameter, and brain, lung and liver tissue was collected for staining.

### Tumor dissociation

Tumors were collected, cut into 1 mm pieces, and incubated in a digestion cocktail of serum-free RPMI supplemented with 1 mg/mL collagenase I (Gibco, #17100017), 250 U/mL collagenase IV (Gibco, #17104019), 20 U/mL DNase I (Invitrogen, #18047019), on a plate rocker at 37^1^C for 60 minutes. Digestion was stopped by adding neutralizing media (1/2 the volume of the digestion reaction), composed of RPMI supplemented with 5% FBS and 2mM EDTA (Invitrogen, #15575020). To obtain a single cell suspension, digested tumors were strained through a 40 um strainer (Falcon, #352340) and washed with neutralizing buffer. Cells were then stained and prepared for analysis by flow cytometry.

### Nanostring

CAR T cells were collected 9 days post-activation, and the CD8+ fraction was isolated using a negative selection kit (Miltenyi Biotec, #130-096-533). Whole cell lysates were generated by vortexing cells for 60 seconds in RLT buffer (Qiagen, #79216) and lysates were stored at -80^1^C. Gene expression analysis used the nCounter CAR-T Characterization Panel, which was supplemented to detect MMPs and OPN, at the Baylor College of Medicine Genomic and RNA Profiling Core using the nCounter Analysis System. Data were analyzed using nSolver 4.0 software (Nanostring).

### Statistical analyses

Two-way ANOVA with Dunnett’s multiple comparisons was used to analyze T cell luminescence in the NB mouse model, mixed-effects analysis with multiple comparisons was used to test Halo infiltration. Unpaired *t*-tests were used to compare tumor luminescence on day 29 in NB model, and HER2 CAR T infiltration into BC tumors. All other data were compared using one-way ANOVA with Tukey’s or Dunnett’s multiple comparisons test. All data were analyzed and graphed using GraphPad Prism (version 10.3.0). Graphs indicate the mean and standard deviation, except where indicated.

---

**Supplementary Figure 1.**
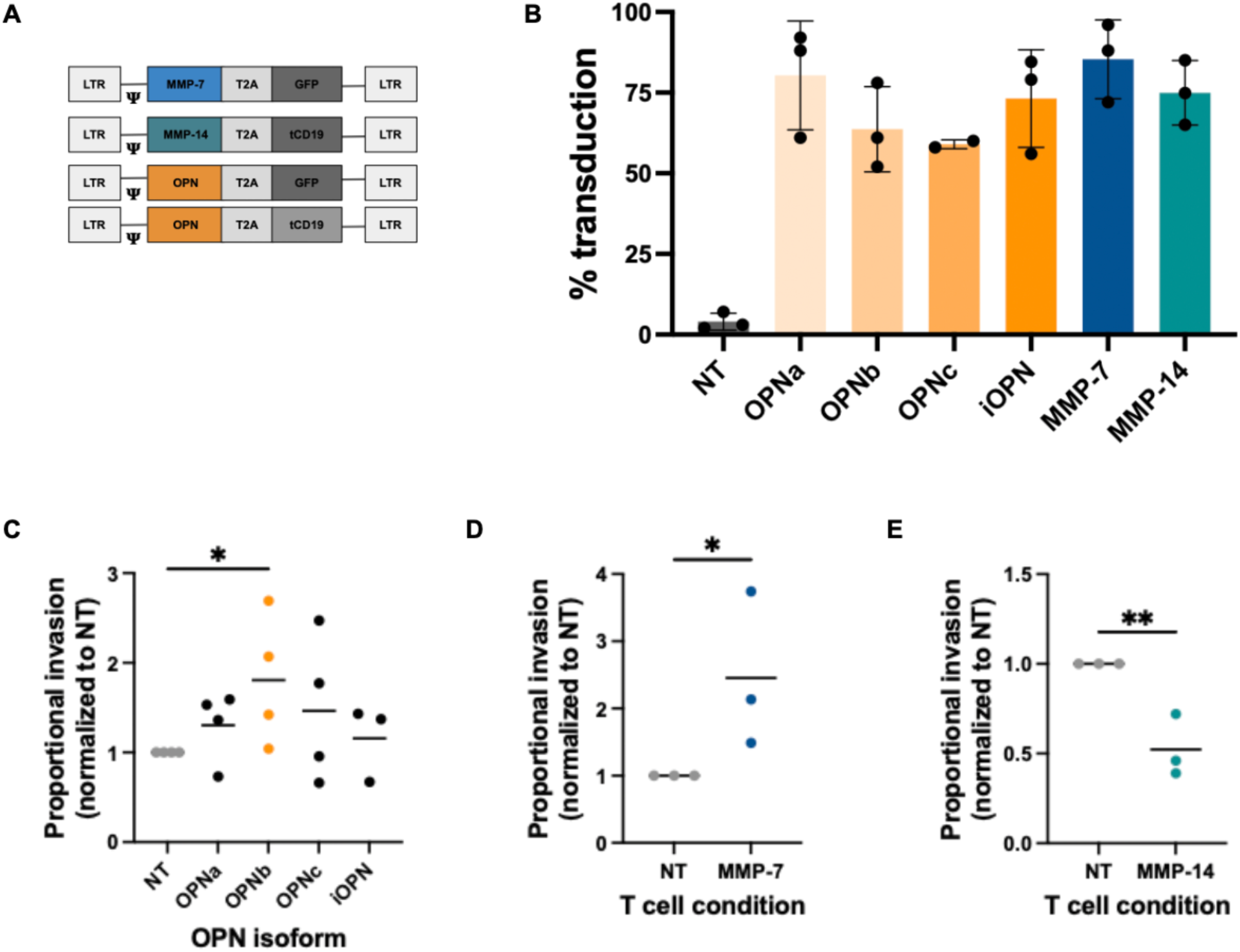
OPN or MMP-7 expression enhances primary T cell movement through ECM. **A.** Schematic of retroviral vector design. Genes of interest followed by a T2A sequence and transduction marker (either GFP or truncated CD19) were cloned into an SFG vector. **B.** Transduction efficiency of primary T cells transduced with candidate genes (mean, SD, *n = 3*). **C-E.** Transwell invasion assays comparing the infiltration of T cells transduced with different isoforms of OPN (OPNa, OPNb, OPNc, or iOPN) **(C)**, MMP-7 **(D),** or MMP-14 **(E)** to NT T cells. Proportional infiltration of gene-modified cells represents a fold-change over NT cells from the same donor (mean, *n = 3-4,* unpaired t test). *\*p < .05*.

**Supplementary Figure 2.**
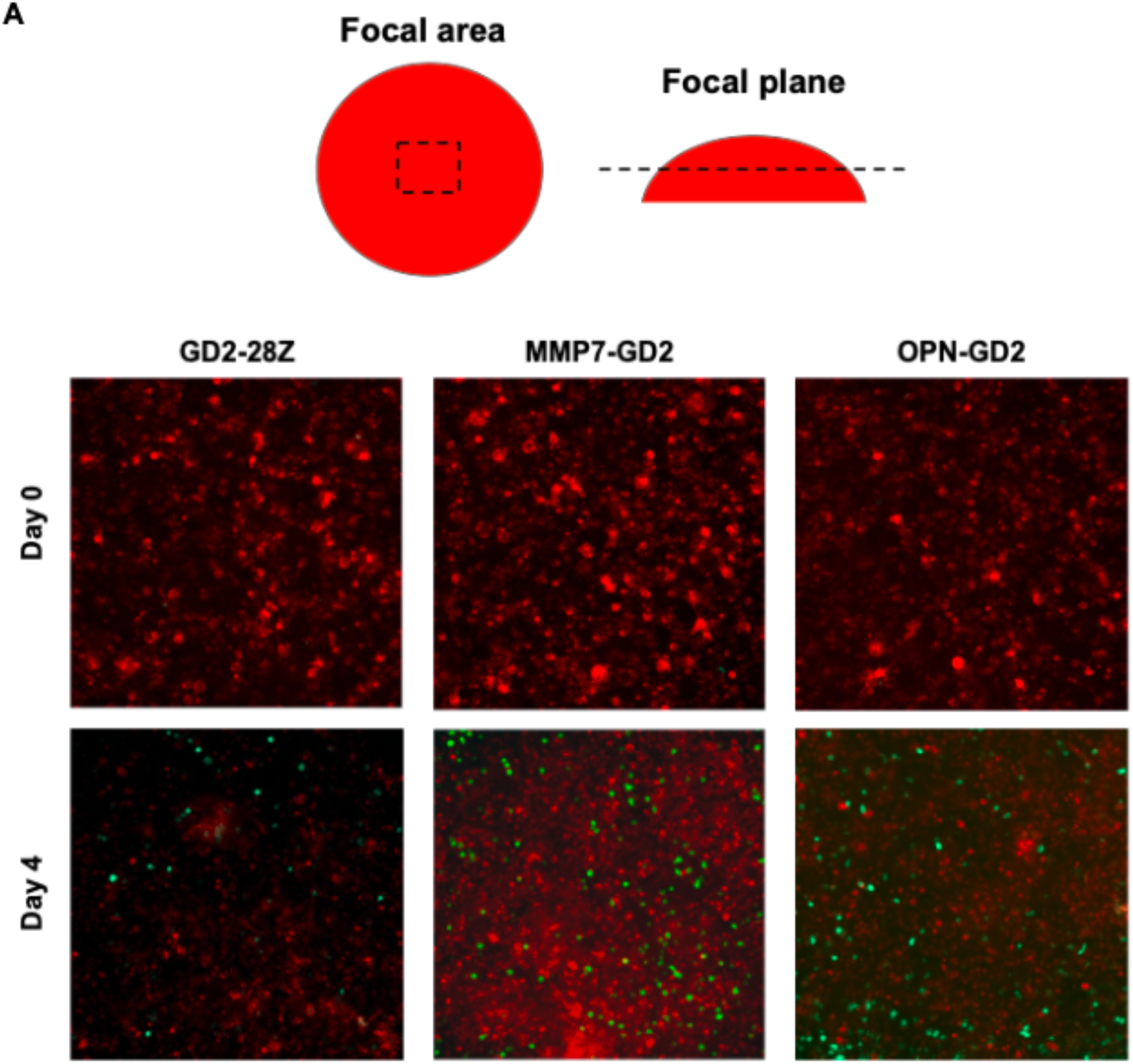
Halo imaging captures infiltration into tumor droplet. **A.** Confocal microscopy images of Halo infiltration assay at t = 5 days with CAR T cells (green) infiltrating central regions of tumor droplets (red). 20X images represent central focal areas and z planes from each droplet.

**Supplementary Figure 3.**
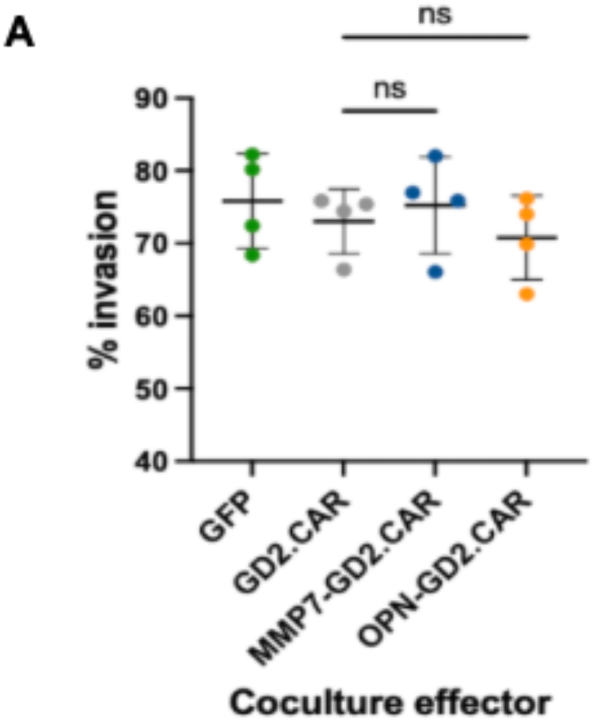
The presence of MMP-7 or OPN does not increase tumor invasion *in vitro*. **A.** Transwell invasion assay comparing the invasion of LAN-1 cells in the presence of supernatants collected from tumor-stimulated GFP, GD2.CAR, MMP7- or OPN-GD2.CAR T cells. Data presented are the mean, SD, *n = 4,* one-way ANOVA with Dunnett’s correction.

**Supplementary Figure 4.**
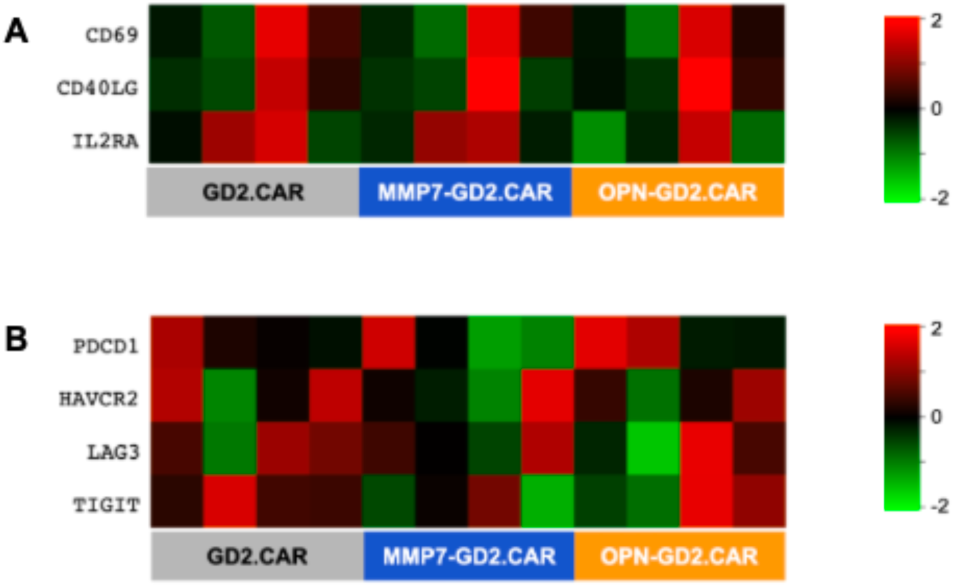
Activation and exhaustion signatures are not dependent on MMP-7 or OPN expression. A-B. Heatmaps of activation (**A**) or exhaustion (**B**) markers in CD8+ GD2.CAR, MMP7-GD2.CAR and OPN.GD2-CAR T cells at rest (day 9 post-activation, *n = 4*).

**Supplementary Figure 5.**
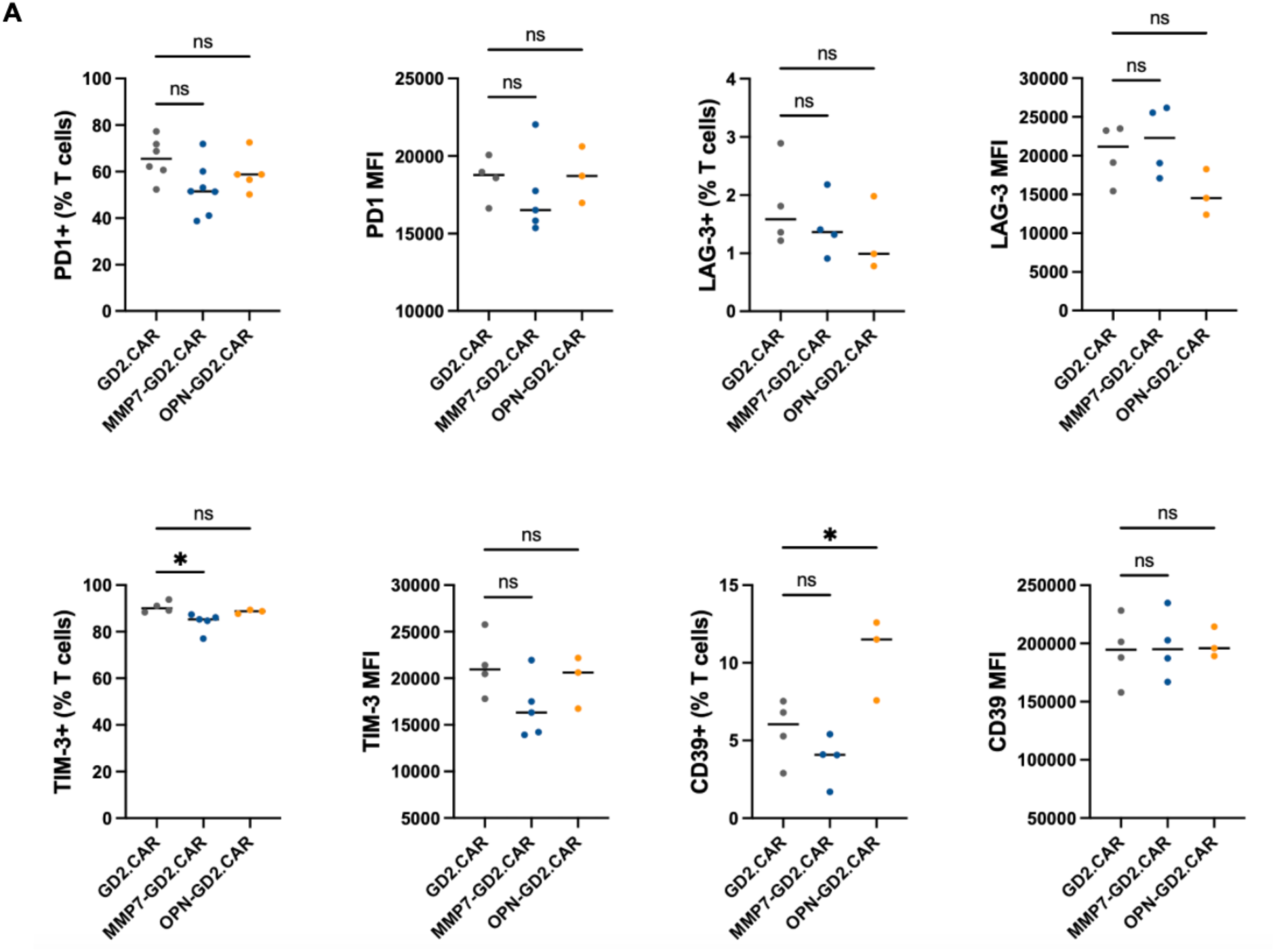
Equivalent expression of exhaustion markers on GD2.CAR and MMP7- GD2.CAR TILs. **A.** Expression of exhaustion markers on CAR T cells collected from dissociated NB tumors ten days post-infusion (median, *n = 2-6*, one-way ANOVA with Dunnett’s correction). \**p < .05*.

## REFERENCES

1. Jackson HJ, Rafiq S, Brentjens RJ. Driving CAR T-cells forward. Nat Rev Clin Oncol. 2016;13(6):370–383. https://www.ncbi.nlm.nih.gov/pmc/articles/PMC5529102/. Accessed Jul 3, 2021. doi: 10.1038/nrclinonc.2016.36.

2. Davila ML, Bouhassira DCG, Park JH, et al. Chimeric antigen receptors for the adoptive T cell therapy of hematologic malignancies. Int J Hematol. 2014;99(4):361–371. Accessed Jul 3, 2021. doi: 10.1007/s12185-013-1479-5.

3. Holstein SA, Lunning MA. CAR T-cell therapy in hematologic malignancies: A voyage in progress. Clin Pharmacol Ther. 2020;107(1):112–122. Accessed Jul 3, 2021. doi: 10.1002/cpt.1674.

4. Richards RM, Sotillo E, Majzner RG. CAR T Cell Therapy for Neuroblastoma. Front Immunol. 2018 Oct 16;9:2380. doi: 10.3389/fimmu.2018.02380. PMID: 30459759; PMCID: PMC6232778.

5. Ahmed N, Brawley V, Hegde M, et al. Autologous HER2 CMV bispecific CAR T cells are safe and demonstrate clinical benefit for glioblastoma in a Phase I trial. J Immunother Cancer. 2015;3(Suppl 2):O11. Published 2015 Nov 4. doi:10.1186/2051-1426-3-S2-O11

6. Haas AR, Tanyi JL, O’Hara MH, Gladney WL, Lacey SF, Torigian DA, Soulen MC, Tian L, McGarvey M, Nelson AM, Farabaugh CS, Moon E, Levine BL, Melenhorst JJ, Plesa G, June CH, Albelda SM, Beatty GL. Phase I Study of Lentiviral-Transduced Chimeric Antigen Receptor-Modified T Cells Recognizing Mesothelin in Advanced Solid Cancers. Mol Ther. 2019 Nov 6;27(11):1919–1929. doi: 10.1016/j.ymthe.2019.07.015. Epub 2019 Jul 30. PMID: 31420241; PMCID: PMC6838875.

7. Park JR, Digiusto DL, Slovak M, et al. Adoptive transfer of chimeric antigen receptor re- directed cytolytic T lymphocyte clones in patients with neuroblastoma. Mol Ther. 2007;15(4):825–833. Accessed Jul 3, 2021. doi: 10.1038/sj.mt.6300104.

8. Feng K, Guo Y, Dai H, Wang Y, Li X, Jia H, Han W. Chimeric antigen receptor-modified T cells for the immunotherapy of patients with EGFR-expressing advanced relapsed/refractory non-small cell lung cancer. Sci. China. Life Sci. 2016;59(5):468–79. doi: 10.1007/s11427-016-5023-8

9. Beatty GL, Haas AR, Maus MV, et al. Mesothelin-specific chimeric antigen receptor mRNA-engineered T cells induce antitumor activity in solid malignancies. Cancer Immunol Res. 2014;2(2):112–120. Accessed Jul 3, 2021.

10. Kershaw MH, Westwood JA, Parker LL, et al. A phase I study on adoptive immunotherapy using gene-modified T cells for ovarian cancer. Clin Cancer Res. 2006;12(20):6106–6115. Accessed Jul 3, 2021.

11. Liu Y, Guo Y, Wu Z, Feng K, Tong C, Wang Y, Hanren D, Shi F, Yang Q, Han W. Anti-EGFR chimeric antigen receptor-modified T cells in metastatic pancreatic carcinoma: A phase I clinical trial. Cytotherapy. 2020;22(10): 573–580.

12. Louis, C. U. et al. Antitumor activity and long-term fate of chimeric antigen receptor-positive T cells in patients with neuroblastoma. Blood 118, 6050–6056 (2011).

13. Heczey, A. et al. CAR T Cells Administered in Combination with Lymphodepletion and PD- 1 Inhibition to Patients with Neuroblastoma. Mol Ther 25, 2214–2224 (2017).

14. Del Bufalo, F. et al. GD2-CART01 for Relapsed or Refractory High-Risk Neuroblastoma. N Engl J Med 388, 1284–1295 (2023).

15. Salmon H, Franciszkiewicz K, Damotte D, et al. Matrix architecture defines the preferential localization and migration of T cells into the stroma of human lung tumors. J Clin Invest. 2012;122(3):899–910. Accessed Jun 21, 2021. doi: 10.1172/JCI45817.

16. Xiao, Z. et al. Desmoplastic stroma restricts T cell extravasation and mediates immune exclusion and immunosuppression in solid tumors. Nat Commun 14, 5110 (2023).

17. Zheng, R. et al. Specific ECM degradation potentiates the antitumor activity of CAR-T cells in solid tumors. Cell Mol Immunol 21, 1491–1504 (2024).

18. Kakarla S, Song X, Gottschalk S. Cancer-associated fibroblasts as targets for immunotherapy. Immunotherapy. 2012 Nov; 4(11): 1129–1138.

19. Winkler J, Abisoye-Ogunniyan A, Metcalf K, Werb Z. Concepts of extracellular matrix remodeling in tumour progression and metastasis. Nat Commun 11, 5120 (2020). doi: 10.1038/s41467-020-18794-x.

20. Henke, E., Nandigama, R. & Ergün, S. Extracellular Matrix in the Tumor Microenvironment and Its Impact on Cancer Therapy. Front Mol Biosci 6, 160 (2020).

21. Rømer, A. M. A., Thorseth, M.-L. & Madsen, D. H. Immune Modulatory Properties of Collagen in Cancer. Front. Immunol. 12, (2021).

22. Murphy K, Weaver C. Janeway’s immunobiology. 9th ed. New York, NY: Garland Science; 2017.

23. Lämmermann T, Bader BL, Monkley SJ, et al. Rapid leukocyte migration by integrin-independent flowing and squeezing. Nature. 2008;453(7191):51-55. Accessed Jul 2, 2021. doi: 10.1038/nature06887.

24. Overstreet MG, Gaylo A, Angermann BR, et al. Inflammation-induced interstitial migration of effector CD4 + T cells is dependent on integrin α V. Nat Immunol. 2013;14(9):949–958. https://www.nature.com/articles/ni.2682. Accessed Jul 2, 2021. doi: 10.1038/ni.2682.

25. Edsparr, K., Basse, P. H., Goldfarb, R. H. & Albertsson, P. Matrix Metalloproteinases in Cytotoxic Lymphocytes Impact on Tumour Infiltration and Immunomodulation. Cancer Microenviron 4, 351–360 (2010).

26. Wolf, K. et al. Physical limits of cell migration: control by ECM space and nuclear deformation and tuning by proteolysis and traction force. J Cell Biol 201, 1069–1084 (2013).

27. Nazha, B., Inal, C. & Owonikoko, T. K. Disialoganglioside GD2 Expression in Solid Tumors and Role as a Target for Cancer Therapy. Front Oncol 10, 1000 (2020).

28. Tadeo, I et al. Extracellular matrix composition defines an ultra-high-risk group of neuroblastoma within the high-risk patient cohort. Br J Cancer 115, 480–489 (2016).

29. Mai, Z., Lin, Y., Lin, P., Zhao, X. & Cui, L. Modulating extracellular matrix stiffness: a strategic approach to boost cancer immunotherapy. Cell Death Dis 15, 1–16 (2024).

30. Caruana, I. et al. Heparanase promotes tumor infiltration and antitumor activity of CAR- redirected T lymphocytes. Nat Med 21, 524–529 (2015).

31. Johnston, A. et al. Engineering self-propelled tumor-infiltrating CAR T cells using synthetic velocity receptors. bioRxiv 2023.12.13.571595. Preprint.

32. Laronha, H. & Caldeira, J. Structure and Function of Human Matrix Metalloproteinases. Cells 9, 1076 (2020).

33. Kaartinen, M. T., Pirhonen, A., Linnala-Kankkunen, A. & Mäenpää, P. H. Cross-linking of Osteopontin by Tissue Transglutaminase Increases Its Collagen Binding Properties*. Journal of Biological Chemistry 274, 1729–1735 (1999).

34. Nishimichi, N. et al. Polymeric Osteopontin Employs Integrin α9β1 as a Receptor and Attracts Neutrophils by Presenting a *de Novo* Binding Site*. Journal of Biological Chemistry 284, 14769–14776 (2009).

35. Weber, G. F., Ashkar, S., Glimcher, M. J. & Cantor, H. Receptor-Ligand Interaction Between CD44 and Osteopontin (Eta-1). Science 271, 509–512 (1996).

36. Zohar, R. et al. Intracellular osteopontin is an integral component of the CD44-ERM complex involved in cell migration. Journal of Cellular Physiology 184, 118–130 (2000).

37. Cancer TODAY, IARC. https://gco.iarc.who.int. Data version: Globocan 2022 version 1.1 (8-2-24).

38. Friedl, P. & Wolf, K. Proteolytic interstitial cell migration: a five-step process. Cancer Metastasis Rev 28, 129–135 (2009).

39. Vicente-Manzanares, M. & Sánchez-Madrid, F. Role of the cytoskeleton during leukocyte responses. Nat Rev Immunol 4, 110–122 (2004).

40. Calderwood, D. A., Shattil, S. J. & Ginsberg, M. H. Integrins and Actin Filaments: Reciprocal Regulation of Cell Adhesion and Signaling *. Journal of Biological Chemistry 275, 22607–22610 (2000).

41. Mori, T. et al. Structural Basis for CD44 Recognition by ERM Proteins. J Biol Chem 283, 29602–29612 (2008).

42. Zhao, H. et al. The role of osteopontin in the progression of solid organ tumour. Cell Death Dis 9, 1–15 (2018).

43. Moorman, H. R. et al. Osteopontin: A Key Regulator of Tumor Progression and Immunomodulation. Cancers (Basel) 12, 3379 (2020).

44. Rittling, S. R., Chen, Y., Feng, F. & Wu, Y. Tumor-derived Osteopontin Is Soluble, Not Matrix Associated *. Journal of Biological Chemistry 277, 9175–9182 (2002).

45. Schreiber, K., Rowley, D. A., Riethmüller, G. & Schreiber, H. Cancer Immunotherapy and Preclinical Studies: Why We Are Not Wasting Our Time with Animal Experiments. Hematology/Oncology Clinics of North America 20, 567–584 (2006).

46. Tong, S., Neboori, H. J., Tran, E. D. & Schmid-Schönbein, G. W. Constitutive expression and enzymatic cleavage of ICAM-1 in the spontaneously hypertensive rat. J Vasc Res 48, 386–396 (2011).

47. Conant, K., Lim, S. T., Randall, B. & Maguire-Zeiss, K. A. Matrix Metalloproteinase Dependent Cleavage of Cell Adhesion Molecules in the Pathogenesis of CNS Dysfunction with HIV and Methamphetamine. Curr HIV Res 10, 384–391 (2012).

